# Glycolipid transfer protein modulates vesicular trafficking from the endoplasmic reticulum in HeLa cells

**DOI:** 10.1101/2025.06.17.657375

**Authors:** Henrik Nurmi, Linda Englund, Alina Henriksson, Max Lönnfors, Peter Mattjus

**Affiliations:** Biochemistry & Cell Biology, Faculty of Science and Engineering, Åbo Akademi University, Artillerigatan 6A, III, 20520 Turku, Finland

**Author notes:** For correspondence: Peter Mattjus.

## Abstract

The glycolipid transfer protein (GLTP) has been proposed to function as a sensor and regulator of glycosphingolipid homeostasis in the cell, as levels of GLTP directly influence the quantities of many glycosphingolipids. Furthermore, through its interaction with the endoplasmic reticulum (ER) membrane protein VAPA (vesicle-associated membrane protein-associated protein A), GLTP may also be involved in regulating intracellular vesicular transport. Here we show that GLTP knockout leads to the sequestration of COPII coat proteins Sec23A and Sec31A at the ERES. The intracellular localization of the small GTPase Sar1A is altered in GLTP knockout cells and in cells expressing a GLTP mutant incapable of VAPA binding, indicating a correlation between the inhibition of the GLTP/VAPA interaction and the altered localization of Sar1A. We also observed alterations in the intracellular localization of the Sar1A-activating GEF Sec12 in GLTP knockout cells, implying a Sec12-activating role of the GLTP/VAPA interaction. Knockout of GLTP does not alter the amounts of other lipid transfer proteins or glucosylceramide synthase, indicating that a decrease in the levels of these proteins is not responsible for the decreased GSL levels associated with GLTP knockout. We propose a role for GLTP in modulating the COPII vesicle trafficking pathway, thereby indirectly regulating the cellular glycosphingolipid homeostasis by controlling the vesicular ceramide transport from the ER to the Golgi. Moreover, the discovery that GLTP localizes to the nucleus at the onset of the DNA-replicating S-phase of the cell cycle introduces an entirely new and unexpected dimension to GLTP’s *in vivo* function.

## Introduction

The glycolipid transfer protein (GLTP) was first discovered in 1980 in lysate from bovine spleen and further characterized in 1982 (1, 2), with its crystal structure being determined in 2004 (3, 4). Protein homologs with similar structural conformations have since been identified, and GLTP is now considered the prototypical member of a superfamily of structurally and functionally similar non-enzymatic lipid transport proteins, termed the GLTP superfamily (5). As its name suggests, GLTP specifically binds glycolipids as ligands (6). However, beyond the requirement that the ligand must be a glycolipid, GLTP exhibits low specificity and can bind most glycolipid and glycosphingolipid (GSL) species (6, 7). GLTP also appears to demonstrate some level of selectivity for acyl chain lengths in its ligands, with shorter chain lengths being slightly more preferred over longer-chain ligands. These characteristics have led to the hypothesis that GLTP may act as a sensor and possibly a regulator of intracellular levels of various glycosphingolipid species (8). In previous work, we have shown that GLTP levels in the cell affect the intracellular levels of numerous species of glycosphingolipids, lending further credence to the sensor-regulator hypothesis (8–10). Specifically, we demonstrated that GLTP knockout (GLTP KO) results in a significant reduction in the intracellular levels of several glycosphingolipid species among those analyzed, indicating that GLTP plays a role in regulating the metabolism of these lipid species (9, 10).

The question of how GLTP regulates GSL homeostasis in the cell is complex, as its known cellular interactions are limited despite its cytosolic localization. GLTP possesses a diphenylalanine-in-an-acidic-tract (FFAT) motif (amino acids 32–38), which enables it to interact with and bind to vesicle-associated membrane protein-associated protein A (VAMP-associated protein A, or VAPA) at the surface of the endoplasmic reticulum (ER) (11). The presence of an FFAT motif is a common feature of lipid transfer proteins (LTPs), with transporters such as CERT, OSBP, Nir2, FAPP2, and ORP3 also featuring FFAT motifs which target them to the ER surface (12–15). These LTPs have also been shown to regulate the intracellular levels of their associated lipids; FAPP2, for instance, regulates the levels of higher-order GSLs derived from glucosylceramide (GlcCer) (16). Being targeted to the ER surface would place GLTP in proximity to the ceramide synthases (17), and by extension the GSL synthesis pathway. However, GLTP requires a carbohydrate headgroup in its ligands to enable binding (4, 6), meaning it can bind complex glycosphingolipid species, whose levels it also regulates, but cannot bind ceramide that lacks a carbohydrate headgroup. Therefore, it is unlikely that GLTP regulates GSL levels by sensing or modulating ceramide levels, especially since not all GSL species, nor sphingomyelin (SM) levels, are affected by GLTP KO (10). We have previously shown that GSL ligand binding diminishes the ability of GLTP to interact with VAPA (8), suggesting that the lipid transport function of GLTP and its ER targeting may be mutually exclusive. Instead, we propose the hypothesis that, while GLTP may not regulate cellular ceramide levels directly, it may influence the vesicular transport of ceramide between the ER and the *cis*-Golgi, thereby contributing to the regulation of GSL homeostasis.

In earlier work, we have shown that the absence of GLTP appears to disrupt the vesicle-based trafficking of membrane-bound proteins from the ER (10). We also demonstrated that this effect could be replicated by expressing a ΔFFAT mutant of GLTP, in which the FFAT motif was altered through point mutation to prevent interaction with the major sperm protein (MSP) domain of VAPA. Cells expressing ΔFFAT-GLTP exhibited similarly disrupted vesicular trafficking as GLTP KO cells (10). This indicates that it is the interaction of GLTP with VAPA at the ER membrane which facilitates the effect that GLTP has on vesicular trafficking in the cell, also indicating that GLTP could act as a regulator of vesicular transport. This, in turn, provides a potential explanation for how GLTP could regulate GSL homeostasis in the cell. Although not yet conclusively proven, it is believed that the pool of ceramide used as the precursor for glycosphingolipid synthesis is primarily transported to the *cis*-Golgi via vesicles originating from the ER (18, 19). This would explain the mechanism through which GLTP would indirectly be able to regulate the GSL levels in the cell, with its ligand binding acting as a regulatory mechanism for vesicular ceramide export from the ER and subsequent synthesis of glycosphingolipids in the Golgi apparatus.

The aim of this study is to characterize the role of GLTP in regulating the vesicle-mediated transport between the ER and the *cis-*Golgi, as well as its role in facilitating the formation of COPII-coated vesicles at the ER surface. Additionally, we aim to characterize the effects of GLTP KO on other lipid transporters and proteins associated with vesicular transport; the knockout of GLTP significantly decreases the cellular levels of GlcCer and GlcCer-derived GSLs (10), but does not completely abolish them, indicating a compensatory mechanism for ceramide transfer between the ER and *cis*-Golgi. The intracellular localization of other lipid transporters is in some cases altered, indicating a possible compensatory role for lipid transfer and/or GSL synthesis of these proteins in the absence of GLTP. We also show that the intracellular distribution of Sar1 is altered in GLTP KO and ΔFFAT-GLTP-expressing cells. The small GTPase Sar1 is responsible for initiating the assembly of the protein coat of COPII vesicles and is activated and recruited to the ER membrane by its guanine-nucleotide exchange factor Sec12 (20–22). The alteration of Sar1 localization aligns with the observed disruption of COPII vesicle traffic and indicates a possible failure of Sec12-mediated activation of Sar1, induced by the absence of the FFAT-motif-mediated GLTP/VAPA interaction. Furthermore, we show that the localization of GLTP changes according to the cell cycle. While GLTP is primarily distributed throughout the cytosol, we observed that the onset of the S phase is accompanied by its translocation to the nucleus.

## Results

### Knockout of GLTP disrupts the COPII vesicle trafficking pathway

GLTP KO has been previously shown to disrupt the transport of GFP-VSVG protein construct from the ER exit sites (ERES) to the plasma membrane. In our previous study, GFP-VSVG was expressed in cells and its transport visualized through fluorescence microscopy, showing that the protein construct was retained at the ERES in GLTP KO cells even two hours after the temperature-induced release of the construct from the ERES, at which time the construct was already beginning to accumulate at the plasma membrane in WT cells. Here, to determine whether GLTP KO genuinely disrupts vesicle transport and budding or if it instead disrupts e.g. cargo protein incorporation, we studied and compared the distribution of COPII vesicle coat proteins in WT and GLTP KO HeLa cells through immunofluorescence imaging. The coat proteins Sec23A and its paralog Sec23B and Sec31 (the two former forming the inner COPII coat layer together with Sec24 and the latter forming the outer layer together with Sec13, respectively (Figure 1A) (23, 24)) were imaged in order to visualize the distribution of COPII vesicles in the cell.

**Figure 1.**
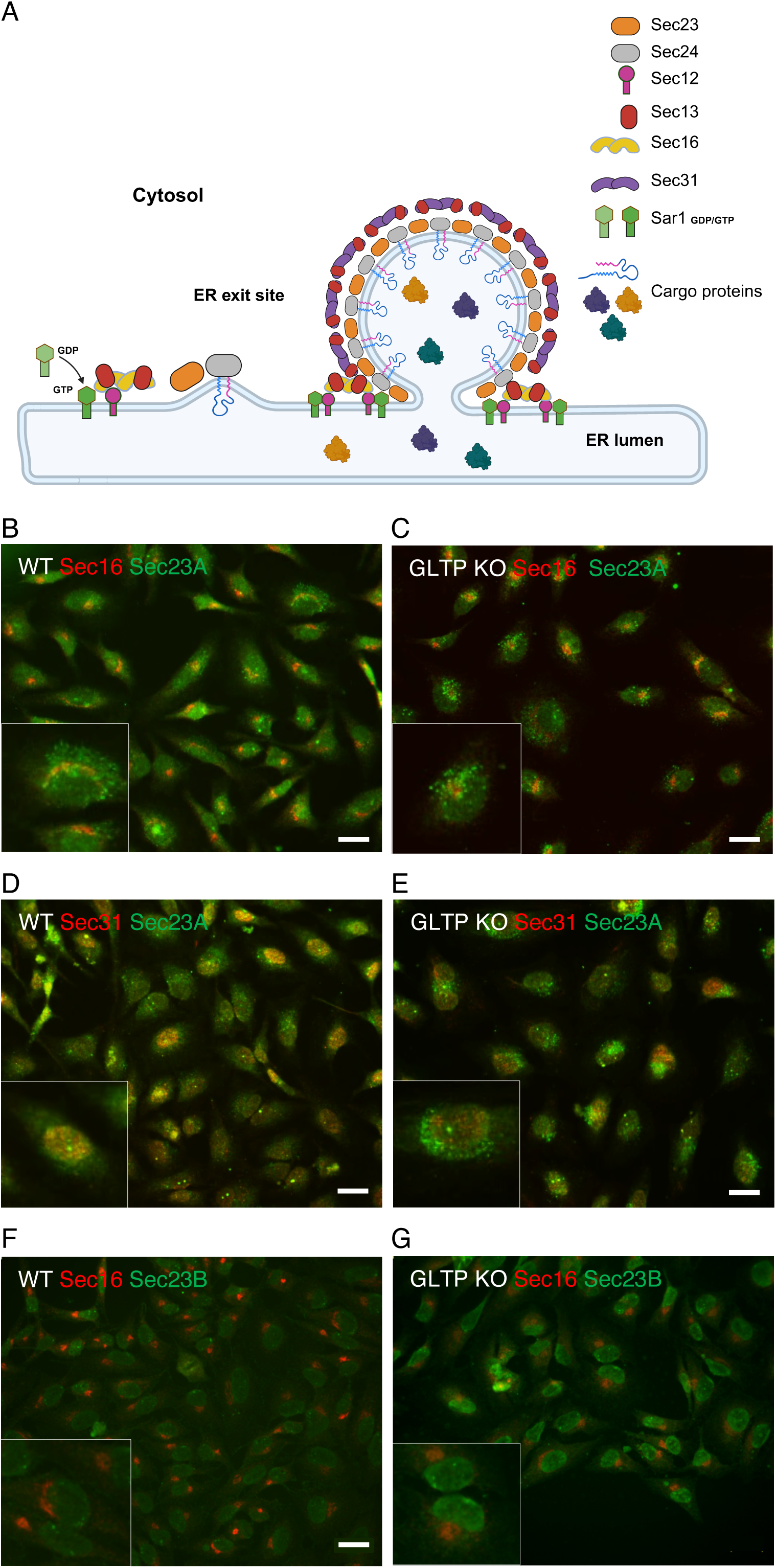
COPII coat proteins Sec23A and Sec31 are retained at the ERES and Sec23B localizes to the nucleus in GLTP KO cells. (A) Schematic presentation of the COPII arrangement at the ERES. Seven proteins build up the COPII coat on the cytosolic side of the ER membrane. The guanine nucleotide exchange factor Sec12 activates Sar1, which inserts into the ER membrane and recruits Sec16-Sec13 and inner heterodimer coat proteins Sec23-Sec24. Outer heterodimer coat proteins Sec13-Sec31 are recruited to complete the coat. (B) Representative fluorescence images of immunostaining of COPII coat proteins and ERES in HeLa cells with anti-Sec23A, anti-Sec31, and anti-Sec16 antibodies. Immunostaining of ERES protein Sec16 (red) and COPII coat protein Sec23A (green) in wild-type HeLa control cells, and (C) GLTP KO HeLa cells. (D) Immunostaining of COPII coat proteins Sec23A (green) and Sec31 (red) in wild-type HeLa control cells and (E) GLTP KO HeLa cells. (F) Immunostaining of COPII coat proteins Sec23B (green) and Sec16 (red) in wild-type HeLa control cells and (G) GLTP KO HeLa cells. The scale bar represents 200 µm. The subpanels are higher magnifications (2x) of representative cells. Figure 1A was created in BioRender. Mattjus, P. (2025) https://BioRender.com/sw45qhe.

The ERES marker protein Sec16 was imaged to provide reference for the origin point of the COPII vesicles versus their intracellular distribution. In the WT HeLa cells, Sec23A and Sec31 are seen distributed throughout the cell with some clustering observed around the ERES marked by Sec16 (Figure 1B & 1D); this clustering is to be expected, as Sec23A and Sec31 would be expected to be found in somewhat higher concentration at the ERES as COPII vesicles are being formed. However, in the GLTP KO cells, Sec23A and Sec31A are only found concentrated at and around the ERES, being almost completely absent from the remainder of the cell body (Figure 1C & 1E). This indicates an abrogation of vesicle release from the ERES brought on by the knockout of GLTP in the cells, supporting our previous observations regarding the disruption of vesicular cargo export from the ER in (10). Interestingly, unlike its paralog Sec23A, Sec23B was not found localized to the ERES in GLTP KO cells, instead localizing to the nucleus in GLTP KO conditions (Figure 1F & 1G). Sec23B has been shown to translocate from the ER/Golgi interface to the nucleus as part of a cellular stress response pathway (25), suggesting adverse conditions in GLTP KO cells. This aligns with previous observations regarding e.g. alterations of cellular metabolism in GLTP KO cells (10).

Representative fluorescence images of immunostaining of Sec23A antibody in HeLa wild-type control cells are shown in Figure 2A and in Figure 2B for GLTP KO HeLa cells. When WT GLTP is reintroduced to the GLTP KO cells, the vesicle release from the ER is restored, with COPII vesicles indicated by their coat protein Sec23A once again being found throughout the cell (Figure 2C). In contrast, expression of the ΔFFAT-GLTP mutant supports our previous findings. Although the mutant retains full glycosphingolipid binding and lipid transport functionality, the mutation in the FFAT motif prevents the interaction of GLTP with VAPA; as a result, COPII coat proteins remain clustered at the ER in these cells (Figure 2D). A quantitative analysis of the Sec23A intensity distribution in the different cell groups is given in the Figure S1. The inability of ΔFFAT-GLTP to interact with both VAPA and VAPB was validated using a pull-down assay using a truncated form of VAP lacking the short C-terminal transmembrane domain, expressed as a GST fusion protein called GST-MSP VAPA (44 kD) or GST-MSP VAPB (40 kD), Figure 2E). The construct includes the GST tag and the cytosolic MSP domain of the VAP protein, which is required for protein-protein interaction; the C-terminal transmembrane domain was omitted due to possible folding concerns. This disruption of vesicle release in cells expressing the VAPA-inhibited ΔFFAT-GLTP further indicates the necessity of GLTP/VAPA interaction for COPII vesicle export to function normally.

**Figure 2.**
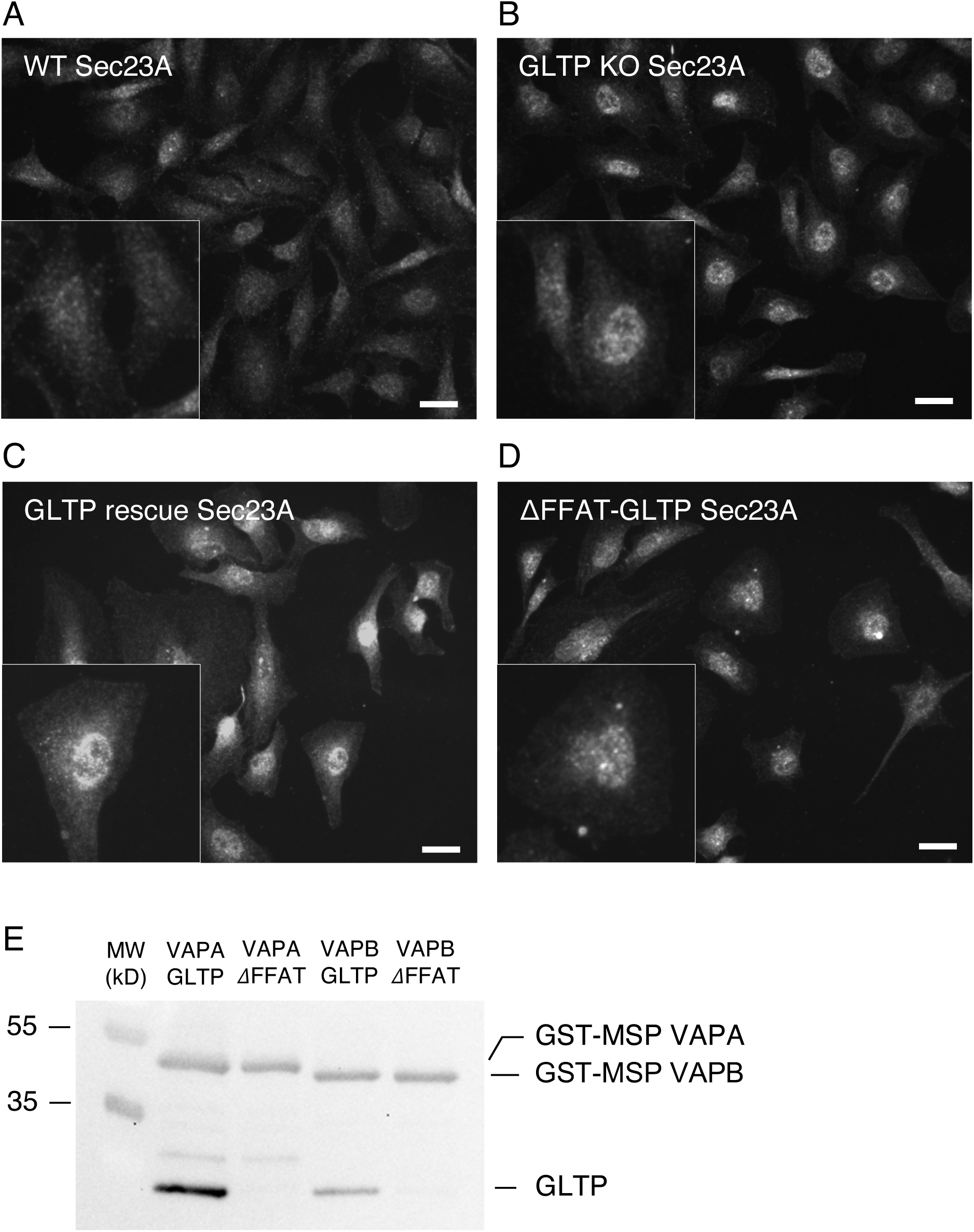
Intracellular localization of Sec23A is restored in GLTP rescue cells but not in ΔFFAT-GLTP cells. Representative fluorescence images of immunostaining of Sec23A antibody staining in (A) HeLa wild-type control cells, (B) GLTP KO HeLa cells, (C) GLTP rescue HeLa cells, (D) and ΔFFAT-GLTP expressing HeLa cells. The scale bars represent 200 µm. (E) Western blot demonstrating GLTP binding to VAPA and VAPB, and the lack of binding by the ΔFFAT-GLTP to VAPA and VAPB. GST-tagged MSP-VAPA and VAPB indicated by anti-GST antibody staining and GLTP indicated by anti-GLTP antibody staining. The scale bar represents 200 µm. The subpanels are higher magnifications (2x) of representative cells.

To further elucidate the mechanism behind the GLTP KO – mediated disruption of COPII vesicle transport, the intracellular localization and levels of the small GTPase Sar1A were assayed. As Sar1A is activated and recruited to the ER membrane by Sec12 to facilitate the assembly of the COPII protein coat, alterations in localization and levels in GLTP KO cells versus WT cells were expected to be observed. Protein levels of Sar1A were not markedly altered in GLTP KO and ΔFFAT-GLTP versus WT and GLTP rescue (Figure 3A). Figure 3B shows the levels of GLTP in wild-type, GLTP KO, GLTP rescue and ΔFFAT-GLTP-expressing cells. Note that the endogenous GLTP band in the wild-type cells is weak due to the strong exposure of the band in the transfected samples. While expression levels of Sar1A were not altered, the intracellular distribution and localization of Sar1 were noticeably affected (Figure 3C-H). In GLTP WT HeLa cells, Sar1A is found in the cytosol and around the nucleus (Figure 3C & 3E); in GLTP KO cells, Sar1A instead appears to withdraw from the cytosol and assemble around the nucleus (Figure 3D), notably distinct and separate from Sec23A (Figure 3D & 3F). In the GLTP rescue and ΔFFAT-GLTP-expressing cells (Figure 3G & 3H) the localization of Sar1A is roughly comparable to that in WT cells and GLTP KO cells, respectively. Although both display a stronger localization of Sar1A towards the nucleus than the WT, the GLTP rescue cells also demonstrate more WT-like cytosolic dispersal of Sar1A (Figure 3E & 3G), whereas in ΔFFAT-GLTP-expressing cells Sar1A appears to be largely sequestered to and around the nucleus, as in GLTP KO cells (Figure 3F & 3H). It becomes apparent that the absence of GLTP-VAPA interaction influences the intracellular distribution of Sar1A and, by extension, COPII vesicle formation and release.

**Figure 3.**
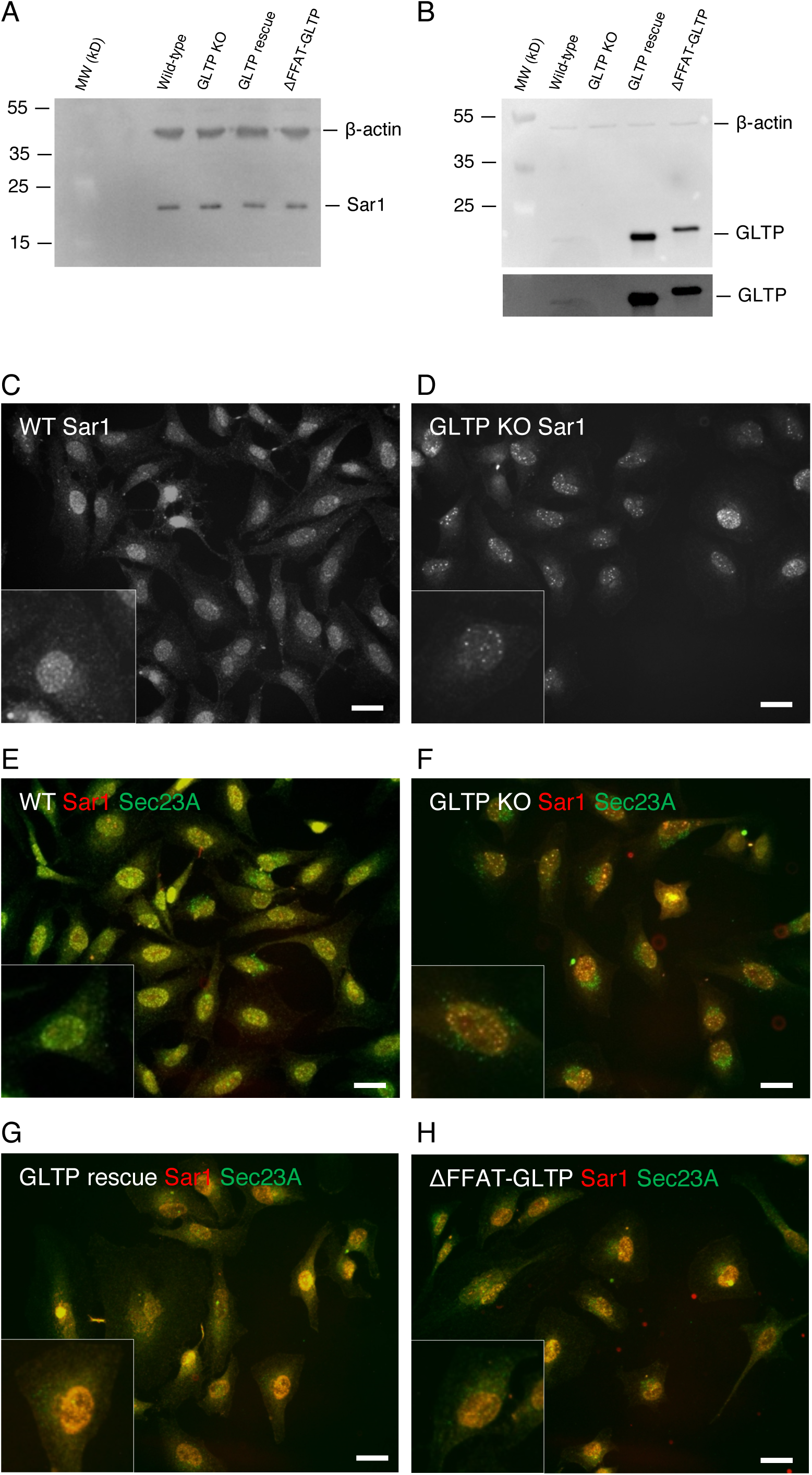
GLTP expression affects localization, but not intracellular levels of Sar1A. (A) Western blot of Sar1A levels in HeLa cell samples: wild-type, GLTP KO, GLTP rescue cells and ΔFFAT-GLTP expressing cells. Beta-actin serves as loading control. (B) Western blot of GLTP levels wild-type, GLTP KO, GLTP rescue cells and ΔFFAT-GLTP expressing cells. The lower panel is a higher exposure of the same blot to visualize the GLTP in the wild-type lane. (C) Representative fluorescence images of immunostaining of Sar1 in HeLa cells stained with anti-Sar1A antibodies WT HeLa cells and (D) GLTP KO cells. (E) Co-immunostaining of Sar1 (red) and Sec23A (green) in WT HeLa cells, (F) GLTP KO cells, (G) GLTP rescue HeLa cells and (H) ΔFFAT-GLTP expressing HeLa cells. The scale bar represents 200 µm. The subpanels are higher magnifications (2x) of representative cells.

### Knockout of GLTP alters the fragmentation and intracellular localization of Sec12

The notable change in the intracellular localization of Sar1A, observed in GLTP KO and ΔFFAT-GLTP expressing cells versus WT HeLa and GLTP rescue cells, appears to indicate either a failure of Sec12 to activate or recruit Sar1A to the ER membrane, or a failure of Sar1 to begin assembling the COPII protein coat once activated. As the only factors required for the assembly of the COPII coat are activated Sar1A and the coat protein complexes themselves (26, 27), it is unlikely that the GLTP/VAPA interaction is involved in the coat formation and vesicle budding once Sar1A has been activated. Therefore, the Sar1A-activating factor, Sec12, was chosen as the next assay target. Sec12 activates Sar1A through the exchange of the Sar1A-bound GDP to GTP, following which Sar1A associates to the ER membrane and sequentially binds and recruits the COPII coat protein complexes. This process catalyzes the GTPase activity of Sar1A, with the GTP hydrolysis deactivating Sar1A and causing its dissociation from the vesicle budding complex (26, 28, 29). With the initial activation and localization of Sar1A to the ERES being reliant on the activity of Sec12, the alteration of its localization in GLTP KO cells implied a potential disruption in Sec12 function in these cells.

As with Sar1A, the cellular expression levels and intracellular localization of Sec12 were assayed through Western blotting and immunofluorescence imaging, respectively. While the expression levels of the canonical Sec12 protein (∼48 kD) were not altered between WT and GLTP KO cells (Figure 4A), certain smaller protein fragments approximately 20 and 25 kD in size, also labeled by the Sec12 antibody, were discovered to be present in WT cell samples and either absent or significantly diminished in GLTP KO cell samples. These protein fragments total a combined size of approximately 45 kD, nearly the size of the canonical Sec12 protein; it is possible that these fragments represent the cytosolic beta-propeller domain of Sec12, which constitutes the majority of the structure of Sec12 and which is responsible for binding to Sar1A and Sec16, as well as to COPII-associated protein scaffolds with TANGO1 and cTAGE5 (30–33). Their presence in WT cells and absence or diminished presence in GLTP KO cells presents an interesting line of questioning; it is possible that the cytosolic domain of Sec12 is cleaved away from the ER surface as the COPII budding complex is dismantled and then further cleaved into two parts. This would certainly explain the absence of these fragments in GLTP KO cell samples, where the COPII vesicle export is disrupted and the COPII budding complex is either never formed or never disassociated from the ER. Another alternative is that the fragments represent isoforms of Sec12, much shorter than the shortest currently known isoforms; isolation and sequencing of these fragments will go a long way to elucidating their identity. Immunofluorescence imaging also showed notable differences in the localization of Sec12 between WT and GLTP KO cells; in GLTP KO cells (Figure 4C, E), Sec12 is notably concentrated to the perinuclear area, in contrast to the far more dispersed dispersal throughout the cell in WT cells (Figure 4B, D). All of this taken together indicates that the non-activation of Sar1A in GLTP KO cells is a consequence of Sec12 itself not being activated; however, the specific activation pathway of Sec12 is still unclear. It would appear, though, that the GLTP/VAPA interaction, or some other interaction mediated by the FFAT-like motif of GLTP, is at least partly responsible for initiating the (or a) pathway which activates Sec12 and facilitates the formation and release of COPII vesicles from the ER.

**Figure 4.**
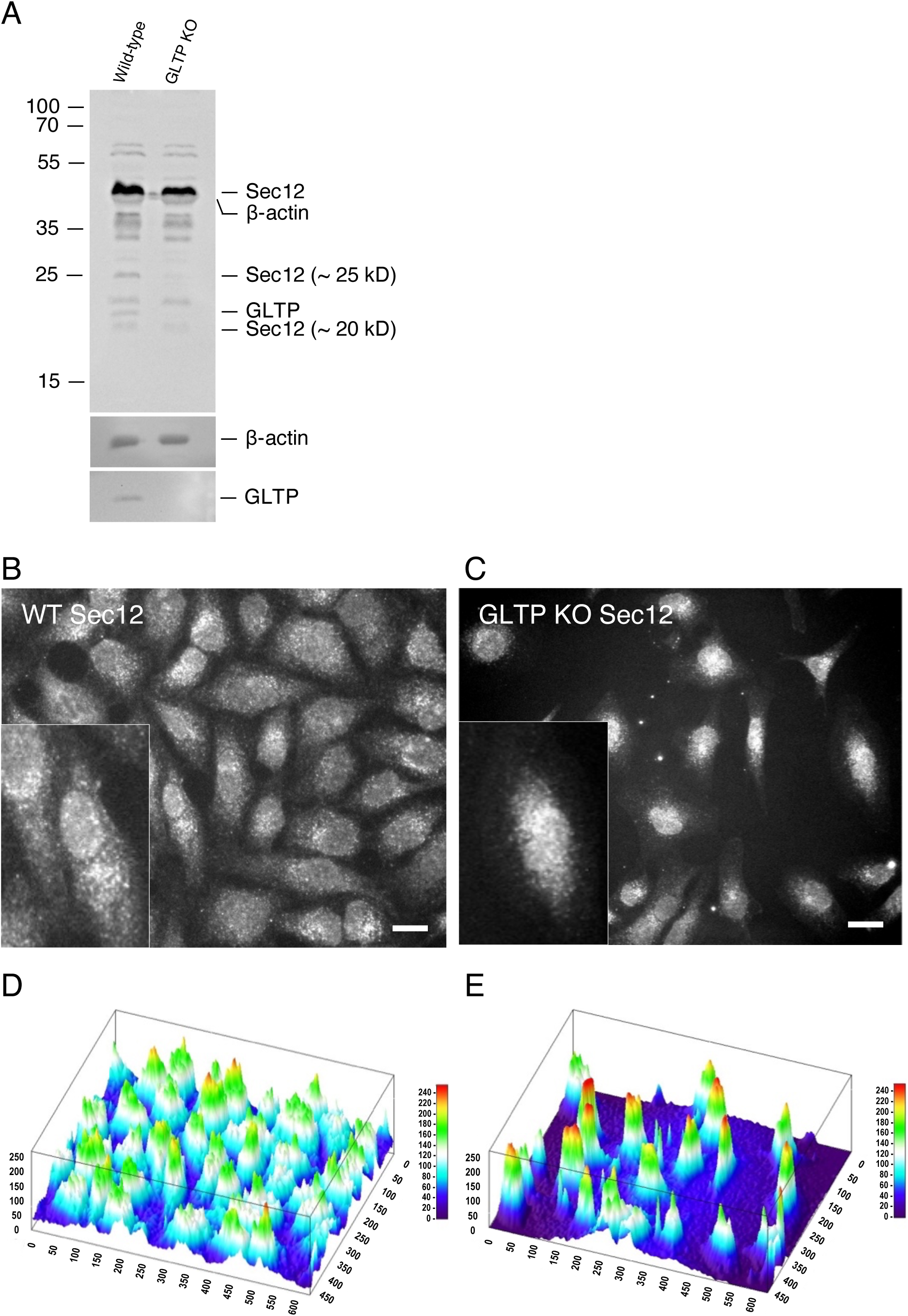
GLTP knockout affects localization and intracellular levels of Sec12. (A) Western blot of Sec12 levels in HeLa wild-type and GLTP KO cells. The full-length Sec12 protein is indicated as Sec12, while the two smaller fragments are labeled Sec12 (∼25 kD) and Sec12 (∼25 kD), respectively. The Sec12 protein bands were detected using an anti-Sec12 antibody and for GLTP an anti-GLTP antibody. Molecular weight markers are shown on the left. Beta-actin serves as loading control. Representative fluorescence images of immunostaining of Sec12 in wild-type HeLa cells (B) and in GLTP KO cells (C). The scale bar represents 200 µm. The subpanels are higher magnifications (2x) of representative cells. Surface plot of the fluorescence intensity of the expression of Sec12 in WT HeLa cells (D) and in GLTP KO HeLa cells (E) visualized using the ImageJ software 3D interactive surface analysis.

### Knockout of GLTP does not alter the cellular levels of other proteins, but does alter their intracellular localization

To study whether GLTP KO affected the levels or intracellular localization of other LTPs or GlcCer synthase (UGCG), protein level analysis through Western blotting and localization analysis via immunofluorescence imaging were carried out on WT and GLTP KO HeLa cells, as well as GLTP rescue and ΔFFAT-GLTP-expressing cells. This allowed us to determine e.g. if other LTPs were expressed in greater amounts or were relocated in the cell to compensate for the lack of GLTP; or if UGCG was decreased because of GLTP KO, possibly contributing to the observed decrease in GSL levels in GLTP KO cells (10). FAPP2, CERT and UGCG were chosen as the primary targets for these assays; FAPP2 and CERT as other LTPs and UGCG due to its role as the key enzyme in the biosynthesis of the GSL precursor GlcCer, catalyzing the GlcCer synthesis using UDP-glucose and ceramide. Sec23A was co-stained alongside the studied LTPs in order to visualize the change occurring in vesicular traffic between WT and GLTP KO cells, as well as to act as a visual reference for the ERES in GLTP KO cells. Golgin-97 was co-stained alongside UGCG as visual reference for the localization of the Golgi apparatus.

The levels of the examined proteins did not change because of GLTP KO (Figure 5A). This indicates that the cell does not compensate for the lack of GLTP by increasing levels of other LTPs as a form of compensatory mechanism; neither did the cellular levels of UGCG change, indicating that the observed decrease in GlcCer levels was not due to e.g. a decrease in synthase levels (Figure 5A). However, as observed through immunofluorescence imaging, the localization of the examined proteins did change even while their levels remained unchanged. As can be observed in the GLTP KO cells, UGCG (Figure 5C) became more concentrated at the ER and around the nucleus compared to the intracellular distribution of UGCG in wild-type HeLa cells (Figure 5B), where UGCG mainly localizes to the Golgi apparatus. We used golgin-97 to provide a reference for the location of the Golgi. In GLTP rescue cells (Figure 5D), the UGCG localization is comparable to the WT HeLa cell UGCG staining (Figure 5B), whereas in the HeLa cells expressing the ΔFFAT-GLTP mutant (Figure 5E) the UGCG localization is comparable to that found in GLTP KO cells (Figure 5C).

**Figure 5.**
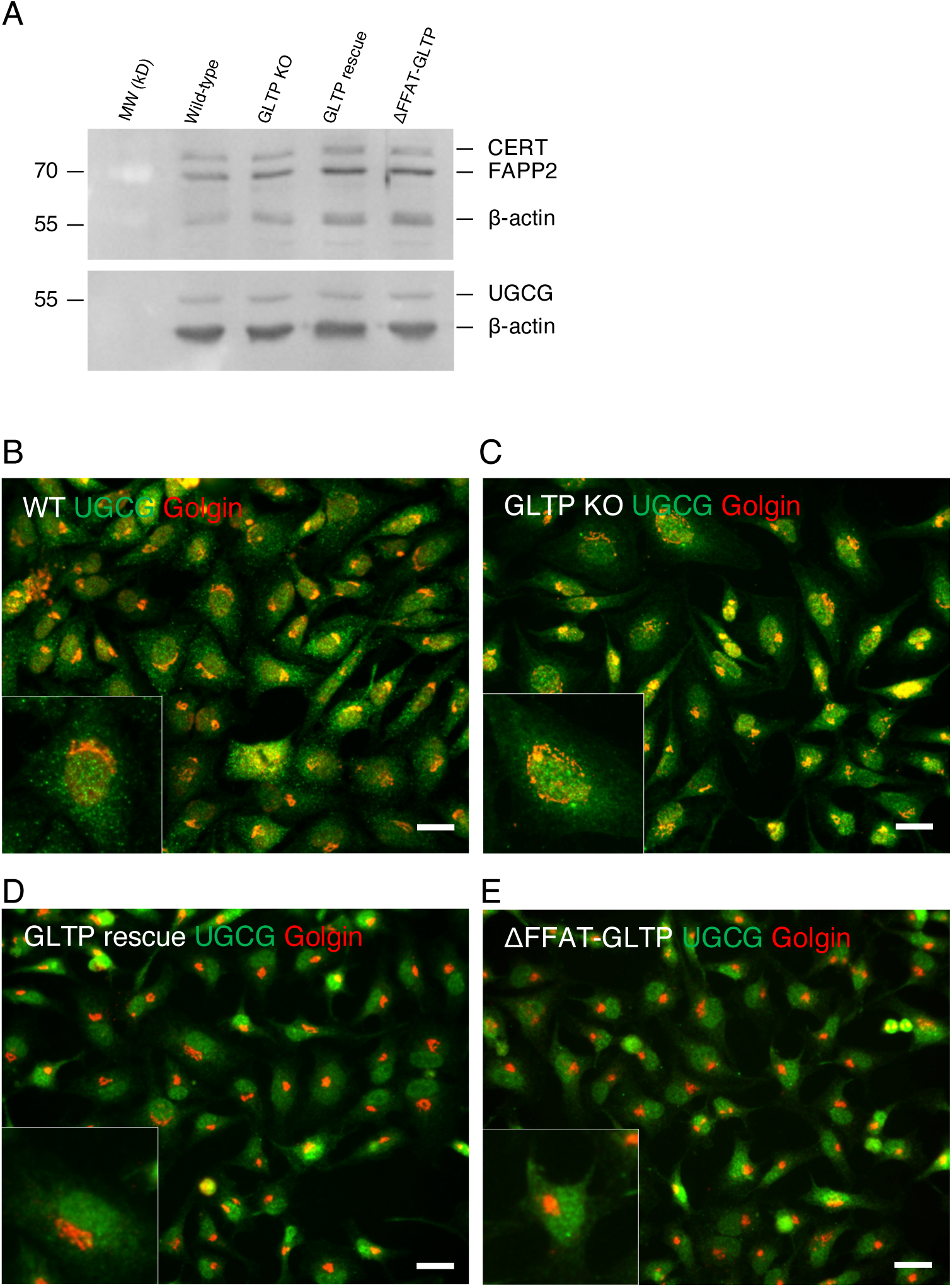
Cells do not compensate for the lack of GLTP by increasing levels of UGCG or LTPs, although protein localizations are altered in the absence of GLTP. (A) Western blot of analysis of the levels of UGCG, FAPP2 and CERT in wild-type HeLa, GLTP KO, GLTP rescue and ΔFFAT-GLTP HeLa cells. Antibodies used are described in the Materials section. Beta-actin was used as a loading control. The protein band in the upper blot that runs between FAPP2, and beta-actin is unidentified. (B) Representative fluorescence images of co-immunostaining of glucosylceramide synthase UGCG (green) and Golgi marker golgin-97 (red) in HeLa wild-type cells and, (C) GLTP KO HeLa cells. (D) Immunostaining of UGCG (green) and golgin-97 (red) in GLTP rescue HeLa cells and (E) ΔFFAT-GLTP expressing HeLa cells. The scale bar represents 200 µm. The subpanels are higher magnifications (2x) of representative cells.

Next, we investigated whether GLTP knockout affects the intracellular localization of other LTPs. In GLTP KO cells, FAPP2 (Figure 6B) and CERT (Figure 7B) both LTPs become more concentrated at the ER, as well as at and around the nucleus as compared to their intracellular distribution in the WT HeLa cells (Figure 6A for FAPP2, Figure 7A for CERT). This change in localization is suggestive of a compensatory role for lipid transport in the absence of GLTP taken on by FAPP2 and CERT; especially FAPP2, as a transporter of GlcCer, is likely to take over the role as primary GSL transporter in the absence of GLTP. In Figure 6E, the FAPP2 localization in GLTP rescue cells is roughly comparable to the FAPP2 localization in wild-type cells (Figure 6A). The lack of FAPP2 concentration towards the nucleus and ER in cells expressing ΔFFAT-GLTP (Figure 6F) supports its compensatory role, as while vesicular trafficking is still disrupted under ΔFFAT-GLTP expression, the ligand-binding and transport functionality of ΔFFAT-GLTP remains intact. ΔFFAT-GLTP therefore likely remains the primary transporter for GSLs in these cells, although GSL synthesis in these cells remains low due to the disruption of ER-Golgi vesicular traffic (10). CERT also becomes more clearly concentrated at the ER and at, as well as around, the nucleus in GLTP KO cells (Figure 7B & 7D) compared to the HeLa wild-type cells (Figure 7A & 7C). A quantitative analysis of the intensity distribution in cells for UGCG, FAPP2 and CERT is given in the Figure S2.

**Figure 6.**
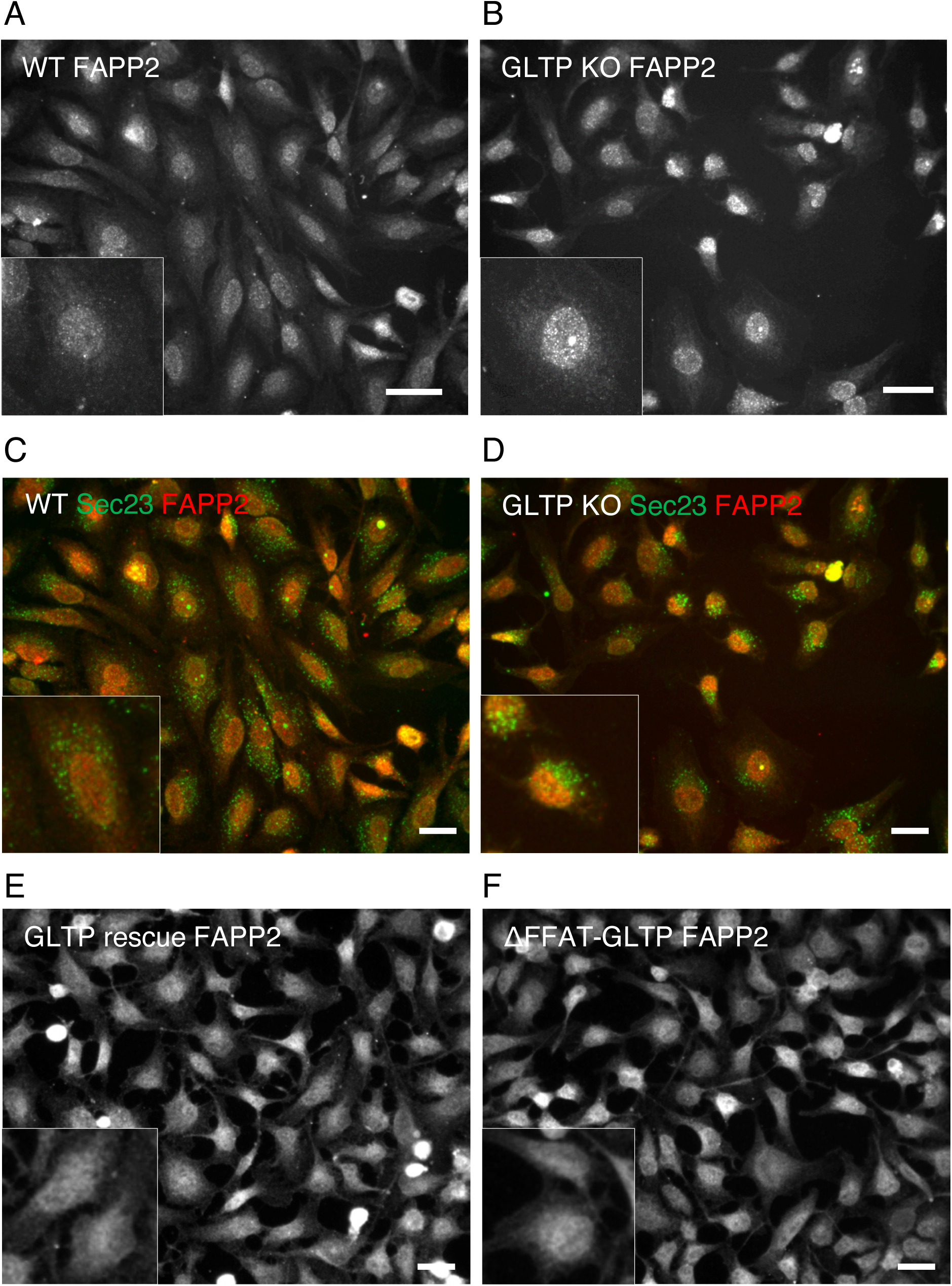
The intracellular localization of FAPP2 is altered in GLTP KO cells. (A) Representative fluorescence images of immunostaining of FAPP2 in wild-type HeLa cells and (B) GLTP KO HeLa cells. Co-immunostaining of FAPP2 (red) and the COPII coat protein Sec23 (green) in, (C) wild-type HeLa cells and, (D) GLTP KO HeLa cells. Immunostaining of FAPP2 in, (E) GLTP rescue HeLa cells and, (F) ΔFFAT-GLTP expressing HeLa cells. The scale bar represents 200 µm. The subpanels are higher magnifications (2x) of representative cells.

**Figure 7.**
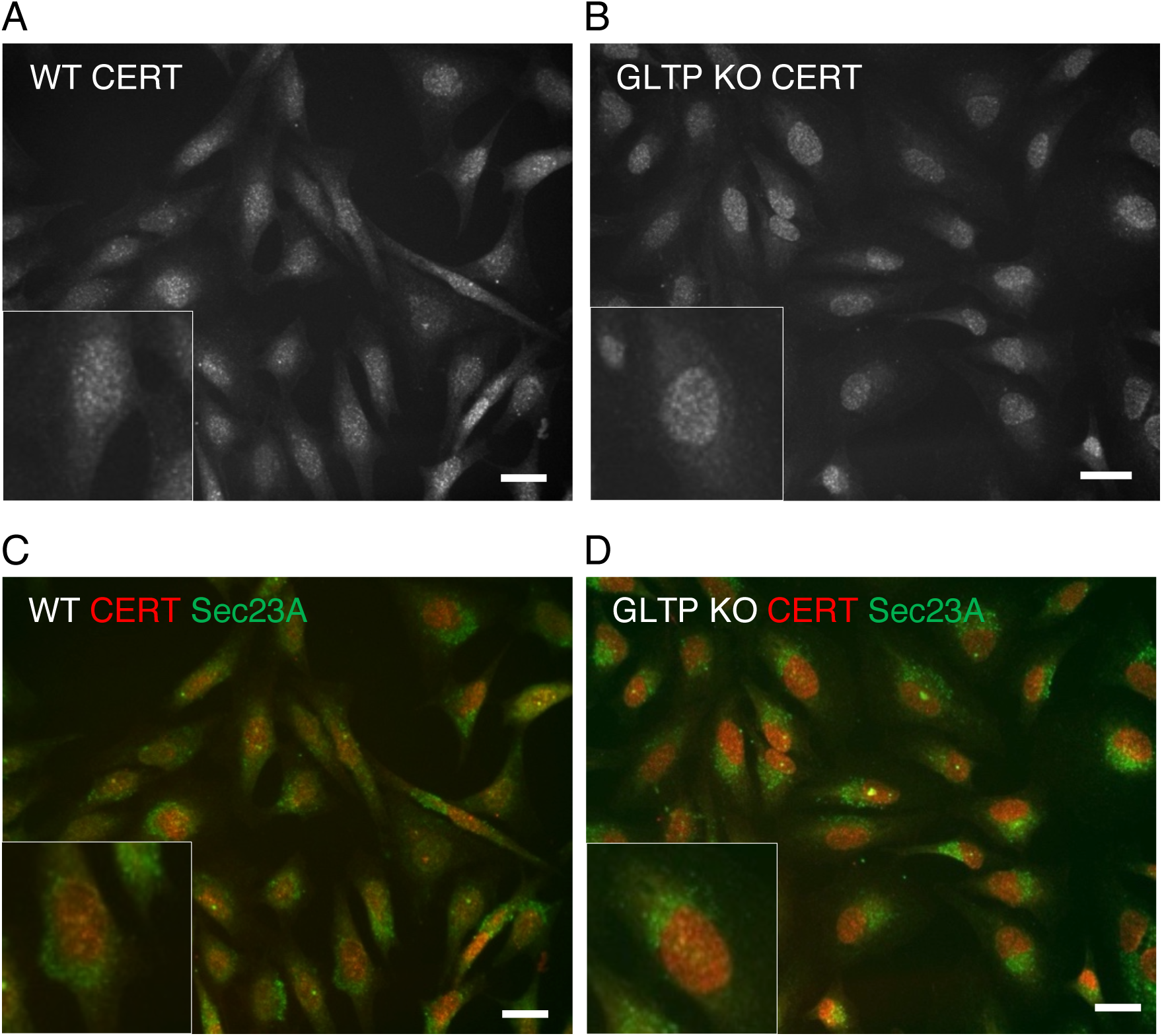
The intracellular localization of the ceramide transporter CERT is altered in GLTP KO cells. (A) Representative fluorescence images of immunostaining of CERT in wild-type HeLa cells and (B) GLTP KO HeLa cells. (C) Co-immunostaining of CERT (red) and the COPII coat protein Sec23A (green) in wild-type HeLa cells and (D) GLTP KO HeLa cells. The scale bar represents 200 µm. The subpanels are higher magnifications (2x) of representative cells.

### Knockout of GLTP alters both cellular levels and intracellular distribution of GSL species

To further study the effect of GLTP KO on intracellular levels of GSLs, an immunofluorescence-based approach was taken to complement the thin-layer chromatography experiments conducted earlier, in which we showed that GLTP knockout significantly reduced cellular levels of multiple glycosphingolipid species, including GlcCer and LacCer as well as both globoside GB_3_ and GB_4_ (10). Commercially available glycolipid antibodies were used to image the localization of glucosylceramide (GlcCer), lactosylceramide (LacCer), and galactosylceramide (GalCer) in WT and GLTP KO HeLa cells. Since GlcCer is synthesized in the Golgi apparatus from ceramide transported there via vesicular transport, and LacCer is subsequently synthesized in the Golgi from GlcCer (18, 34), their localizations were expected to be affected. GalCer, on the other hand, is synthesized in the ER (18), and was therefore not expected to be affected by GLTP KO. As seen in Figure 8, the immunostaining with antibodies towards both GlcCer and LacCer are indeed different in GLTP KO cells compared to the staining in WT HeLa cells, with the effect of GlcCer being especially striking. Both lipids are distributed throughout the cell in WT cells, whereas in GLTP KO cells both GlcCer (Figure 8A & 8B) and LacCer (Figure 8C & 8D) are found more near the ER and nucleus. The increase in fluorescence intensity for GlcCer staining in GLTP KO cells versus WT cells should not be taken as indicative of increased GlcCer levels but rather of the concentration of existing GlcCer towards the ER and nucleus, as our previous quantitative lipidome analysis (10) has in fact shown a significant decrease in GlcCer in GLTP KO conditions. Interestingly, the localization of GalCer was also affected, as can be observed in Figure 8E & 8F. GalCer in GLTP KO cells appears to cluster at certain points close to the nucleus, likely to be the ERES or similar region – co-immunostaining of GalCer and Sec16 at the ERES was unfeasible as both antibodies were rabbit-derived. This indicates that while the levels of GalCer may not be affected by GLTP KO, GalCer is retained in the ER after synthesis due to the lack of an exporter protein – namely, GLTP.

**Figure 8.**
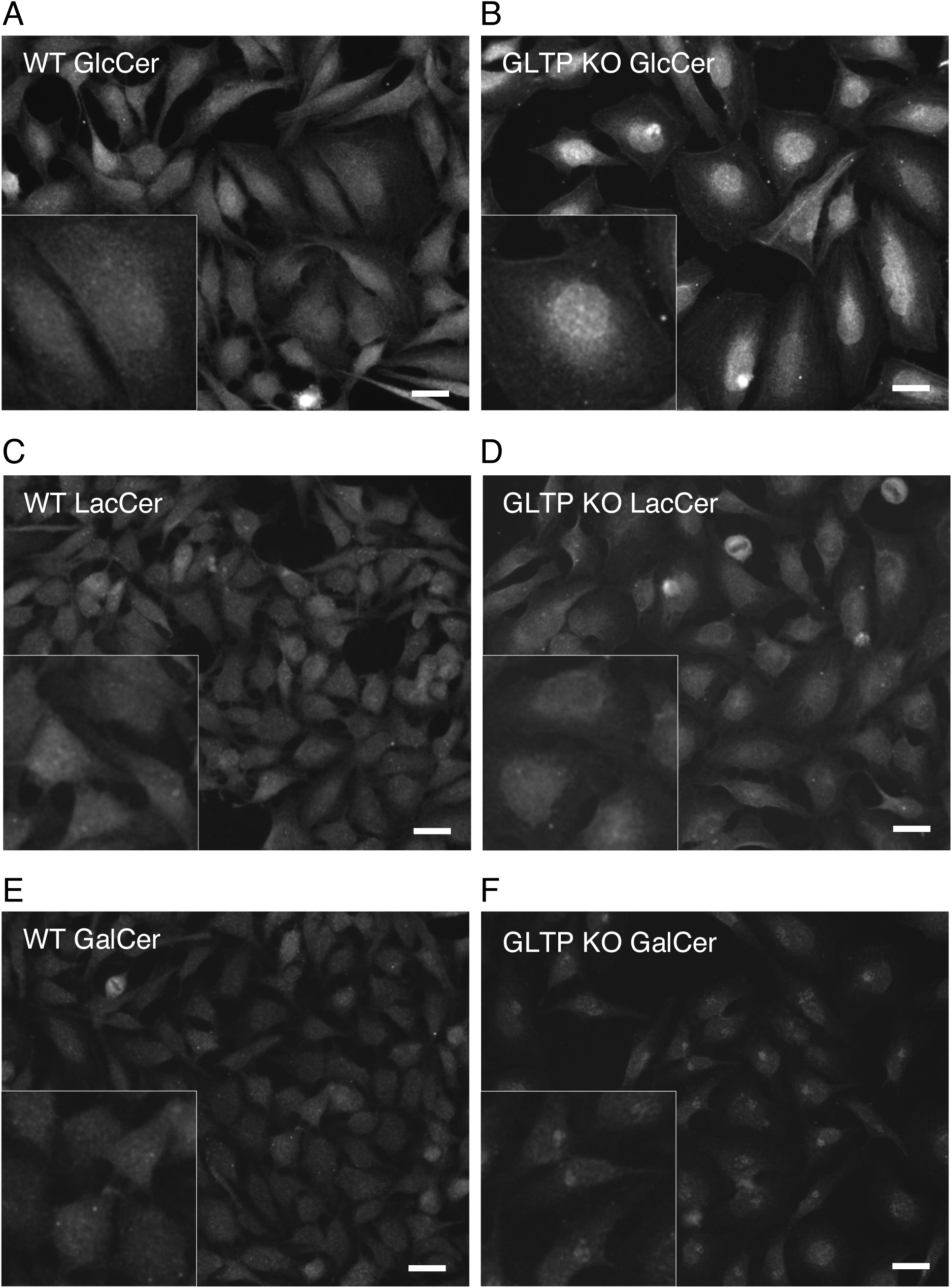
Immunostaining of HeLa cells with antibodies towards GlcCer, LacCer and GalCer are different in GLTP KO cells compared to the staining in WT cells. Representative fluorescence images of immunostaining of glycolipids in wild-type HeLa cells and GLTP KO HeLa cells. (A) Immunostaining of glucosylceramide (GlcCer) in wild-type HeLa cells and (B) GLTP KO HeLa cells. (C) Immunostaining of lactosylceramide (LacCer) in wild-type HeLa cells and (D) GLTP KO HeLa cells. (E) Immunostaining of galactosylceramide (GalCer) in wild-type HeLa cells and (F) GLTP KO HeLa cells. The scale bar represents 200 µm.

Although GLTP knockout significantly reduces cellular GlcCer and its derivatives (10), it does not eliminate them entirely, suggesting the presence of compensatory mechanisms. These may include alternative ceramide transport routes from the ER to the cis-Golgi or a relocation of the GlcCer synthesis machinery. In GLTP KO cells, UGCG appears to shift from the Golgi to perinuclear and ER regions (Figure 5), where GlcCer and partially LacCer also accumulate (Figure 8). This suggests that impaired ER-to-Golgi vesicular transport may cause GlcCer synthesis to occur directly in the ER, with UGCG utilizing ER-localized ceramide. This could explain the persistence of GSLs despite disrupted trafficking, albeit at reduced levels, possibly due to less efficient synthesis in the ER and competition for ceramide with CERT and ceramide galactosyltransferase. Alternatively, GlcCer accumulation in the ER/perinuclear region may result from FAPP2-mediated retrograde transport becoming the dominant pathway in the absence of GLTP (16, 35).

While GalCer synthesis and levels, unlike those of GlcCer derivatives, may be unaffected by GLTP KO, the absence of GLTP still disrupts the export of GalCer from the ER. The disruption of GalCer export would also likely result in a decrease in sulfatide levels corresponding to the decrease in GlcCer and its derivatives, as GalCer is exported to the Golgi for conversion into sulfatide by cerebroside sulfotransferase (CST) in the lumen of the Golgi (36). This would also likely mean that GLTP depletion would have severe consequences for e.g. neuronal development. Mice with conditional knockout of GLTP in their oligodendrocytes have been recently generated, with these mice demonstrating dys- and hypomyelination as well as ER pathologies in their oligodendrocytes; constitutive whole-body GLTP KO proved to be embryonically fatal (37). The GLTP KO – related depletion of GlcCer, in combination with a likely depletion of sulfatide and inability to efficiently transport and utilize GalCer, is likely at least partially the cause of GLTP depletion being fatal to the organism.

### Synchronization assays reveal localization of GLTP to nucleus in S phase of the cell cycle

To determine if the intracellular localization or levels of GLTP changed during the cell cycle, and if the localization of GLTP was associated with any specific phase of the cell cycle, several synchronization assays were carried out. Protein analysis of the cells at the various timepoints for all the synchronizations showed that GLTP levels remained the same throughout the cell cycle (Figure 9A). A double thymidine block was used to arrest cells at the G_1_/S phase boundary by inhibiting deoxynucleotide metabolism and thereby DNA synthesis (38). RO3306 was used to arrest cells at the G_2_/M boundary just before mitosis, by inhibiting CDK1 (39). Nocodazole was used to arrest the cells during mitosis by inhibiting the formation of the mitotic spindle. Immunofluorescence imaging was then used to determine differences in GLTP localization as an effect of the cell cycle; cells synchronized via thymidine block were stained against GLTP and imaged at 180-minute intervals post block release, at 0 minutes, 180 minutes, and 360 minutes. To determine the success of the synchronization and the cell cycle stage at each harvest time point, the cells were analyzed using flow cytometry. In the FACS analysis, the unsynchronized cells (WT and GLTP KO) were used as controls. Cells were synchronized with thymidine and subsequently harvested at 120-minute intervals, at 0 min, 120 min, and 240 min. Since the G_1_ phase is the longest, it is not uncommon for most cells in a population to be in this phase. Here, the control groups showed the highest proportion of cells in the G_1_ phase, with 68.7% of WT cells and 66.4% of KO cells. For the thymidine-treated cells harvested at 0 min, the number of cells in the G_1_ phase decreased and increased in the S phase, where 60.1% of the cells were located. Of the cells harvested 120 min after release from the thymidine block, 78.2% were in the S phase, while 76% of the cells harvested after 240 min were in the S phase (Figure 9B).

**Figure 9.**
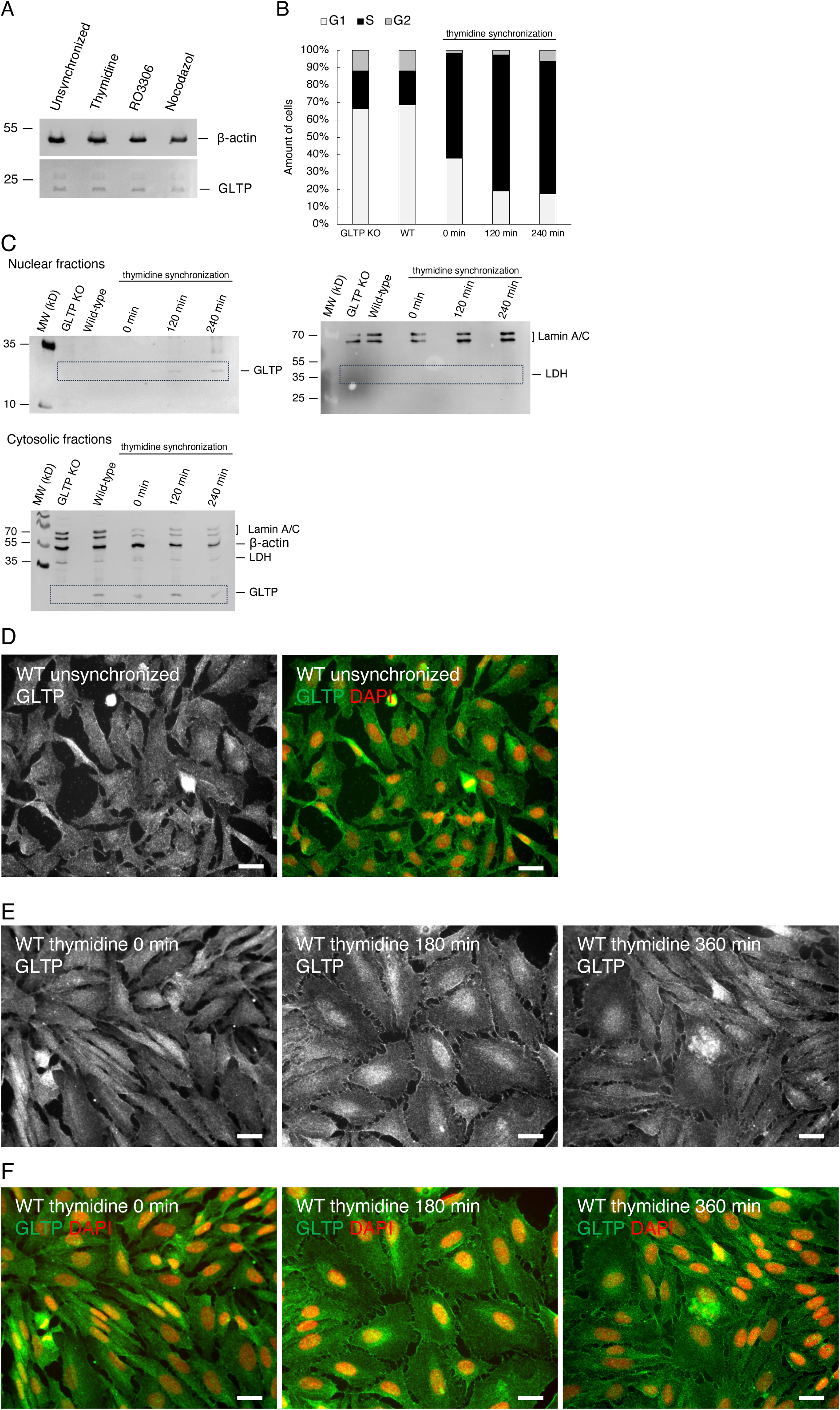
The immunofluorescence imaging of thymidine-synchronized HeLa cells shows a distinct localization of GLTP to the nuclear area of the cells. (A) Western blot analysis of the GLTP protein levels of unsynchronized and cells synchronized with thymidine, RO3306 and nocodazole. Beta-actin was used as a loading control. (B) The percentage of unsynchronized HeLa cells (WT and GLTP KO) and double thymidine (0 min, 120 min, and 240 min) was 60.1%, 78.2%, and 76.0% into the S-phase, respectively. The control cells showed most cells in the G1 phase (68.7% in WT and 66.4% in KO). (C) Western blot analysis at different timepoints for thymidine synchronized cells, (left) shows the nuclear localization of GLTP using anti-GLTP antibodies, (right) the presence of the nuclear marker lamin A/C and the absence of lactate dehydrogenase (LDH) in the nuclear fractions of GLTP KO, WT and the thymidine synchronized cells. The lower blot shows the cytosolic fractions for the double thymidine synchronization samples and the location of LDH and GLTP. (D) Representative immunofluorescence image of GLTP in unsynchronized wild-type HeLa cells. (D) Representative immunofluorescence images of GLTP in wild-type HeLa cells synchronized with thymidine. (E) Immunostaining of GLTP (green) shown co-stained with DAPI (red) as a nuclear marker imaged at different timepoints post release from thymidine block. The scale bar represents 200 µm. Figure 9A created in BioRender. Mattjus, P. (2025) https://BioRender.com/ze4x93c.

Thymidine-synchronized cells were then harvested for protein analysis and the nuclear fraction purified from the lysate. GLTP was found to be present in the purified nuclear fraction at the different timepoints post-thymidine release (Figure 9C, left panel). The presence of the nuclear marker lamin A/C and the absence of lactate dehydrogenase (LDH) in the nuclear fractions of WT, GLTP KO, and the thymidine synchronized cells confirms that there is no contamination of the cytosolic fraction in the nuclear isolate (Figure 9C, right panel).

Compared to unsynchronized wild-type HeLa cells (Figure 9D), immunofluorescence imaging of the thymidine synchronization showed a distinct localization of GLTP to the nuclear area of the cells, with the most evident nuclear localization of GLTP occurring at around the 180 min timepoint following the release of the cells from the thymidine arrest (Figure 9E & 9F).

The localization of GLTP to the nucleus during the DNA-replicating S-phase presents an entirely new potential set of functions for GLTP and an interesting new line of research; while lipids have been proposed to possess a variety of roles in DNA replication and other lipid transfer proteins (e.g. PLTP) have been shown to localize to the nucleus (40, 41), the functions of LTPs in the nucleus have generally not been extensively studied. The mechanism through which the cytosolic GLTP, possessing no known nuclear targeting motif or domain, is localized to the nucleus is thus far unknown, as are its potential functions within the nucleus; however, given the proposed possible functions of lipids in modulating DNA replication and repair and that GLTP localizes to the nucleus during the DNA-replicating S phase, it is possible that GLTP is also involved in DNA replication.

We did not observe changes in the GLTP localization after the treatment with RO3306 (Figure 10A); however, treatment with nocodazole (Figure 10B) also showed potential GLTP localization to the nucleus. If GLTP possesses roles relating to DNA replication, maintenance, and repair, this could explain the localization of GLTP to the nucleus also in association with mitosis; however, this study focused more on GLTP in the S phase as the nuclear localization after thymidine synchronization was more pronounced and longer-lasting than the one seen after nocodazole synchronization.

**Figure 10.**
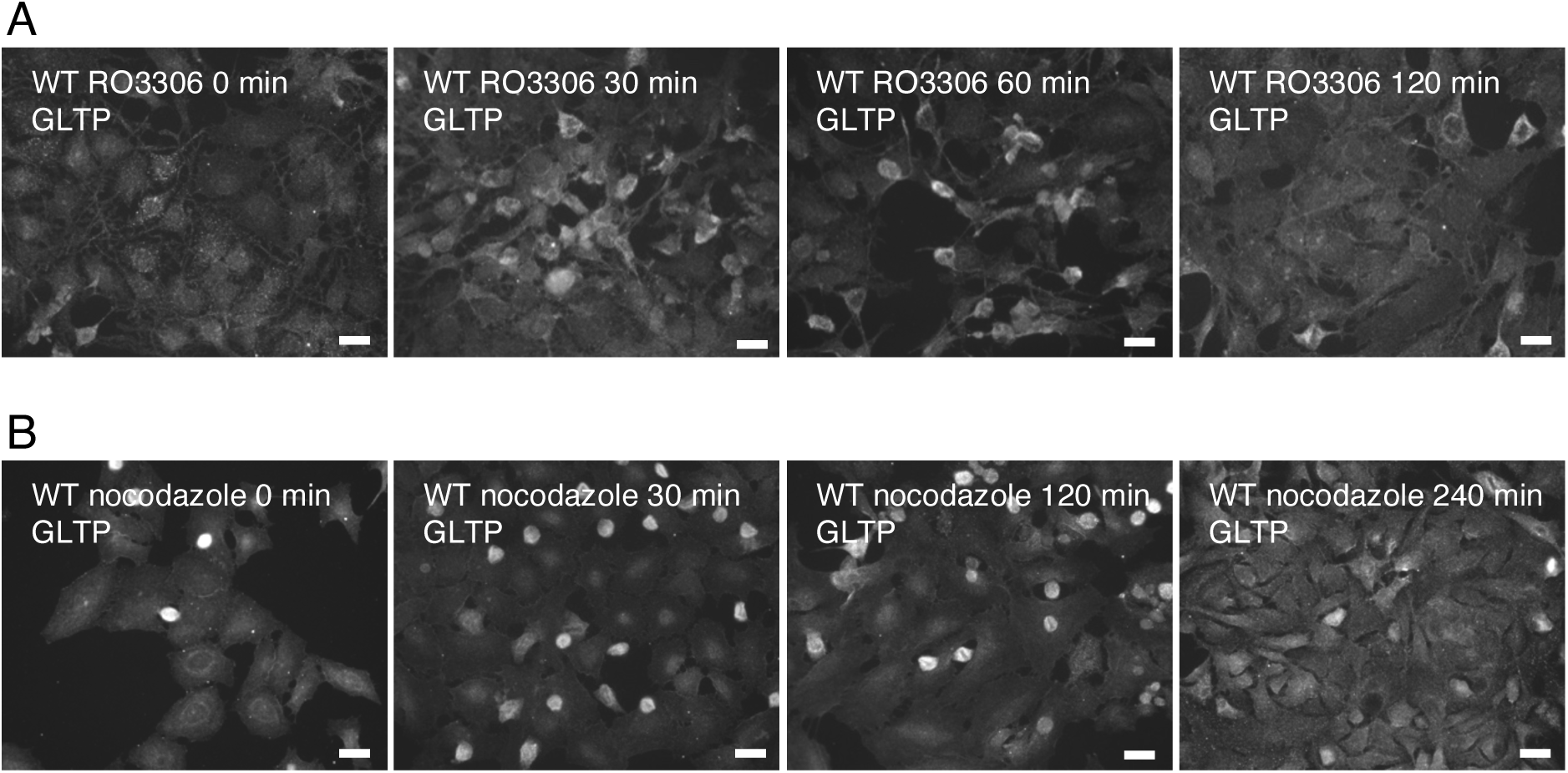
Immunofluorescence imaging of the RO3306 and nocodazole synchronizations and the timepoints thereafter did not show any notable differences in GLTP intracellular distribution as compared to unsynchronized cells. (A) Immunofluorescence images of GLTP in HeLa WT cells synchronized with RO3306 at different timepoints after post release. (B) Immunofluorescence images of GLTP in HeLa WT cells synchronized with nocodazole at different timepoints after post release. The scale bar represents 200 µm.

## Discussion

We have previously shown that GLTP knockout leads to a decrease in the levels of certain glycosphingolipid species in cells, suggesting a regulatory role for GLTP in maintaining GSL homeostasis (10, 42). We also observed that GLTP knockout disrupts the transport of COPII-vesicle-bound cargo from the ER to the plasma membrane. To determine whether the disruption was indicative of e.g. a cargo incorporation defect at the ERES or indeed a disruption of vesicular transport, we observed the behavior of the COPII coat proteins Sec23A and Sec31A to draw more conclusions as to the behavior of the vesicles themselves rather than just their cargo. The retention of Sec23A and Sec31A at the ERES in GLTP KO cells provides ample indication that the absence of GLTP in fact does abrogate at least the release of COPII vesicles from the ER, if not the formation of the vesicles altogether. We further observed that this effect is specifically due to the loss of FFAT-motif-mediated interaction between GLTP and VAPA. Expression of a ΔFFAT mutant GLTP that is structurally and functionally identical to wild-type GLTP except for a point mutation that disrupts FFAT-mediated binding, resulted in similar sequestration of Sec23A and Sec31A at ERES, indicating a failure in vesicle release from the ER. This finding aligns with previous observations that even a protein fragment containing the FFAT motif is sufficient to facilitate vesicle-mediated protein export from the ER (43).

Through assaying the behavior of the small GTPase Sar1A and the guanine nucleotide exchange factor Sec12 in response to the presence or absence of GLTP and GLTP/VAPA interaction, we have also begun to elucidate the mechanism through which the FFAT-motif-mediated interactions of GLTP facilitate COPII vesicle trafficking. While it is well established that Sec12 activates and recruits Sar1A to begin the formation of COPII vesicles, it is less clear how Sec12 itself is activated. Recent studies have suggested different mechanisms as being involved in Sec12 activation, including the recruitment of Sec12 to protein complexes with TANGO1 and cTAGE5 (31), potassium ion binding to Sec12 (44), and the activation through phosphorylation of Sec12 by the ER-resident protein leukocyte tyrosine kinase (LTK) (45). It is plausible that the interaction of GLTP/VAPA and the activation of Sec12 are multiple intermediary steps removed from one another and only one pathway among many possible. Nonetheless, we observe a clear correlation between the inhibition of FFAT-motif-mediated GLTP/VAPA interaction and both the altered intracellular localization of Sar1A and Sec12, as well as the disruption of COPII vesicle secretion from the ER, implying a Sec12-activating role of the GLTP/VAPA interaction.

As the GLTP/VAPA interaction appears to be an essential facilitator of the COPII vesicular export pathway, the observed decreases in the levels of GSLs in GLTP KO cells receive an explanation. As the affected GSL species are all glucosylceramide and derivatives thereof, this indicates a failure either at or before GlcCer synthesis (46). The most reasonable explanation, for which evidence is also here observed and reported, is that the depletion of GlcCer and its derivatives in GLTP KO cells is due to the lack of ceramide transport to the *cis*-Golgi via COPII vesicles, as GlcCer and its derivatives are ultimately all derived from ceramide transported to the *cis*-Golgi via vesicular export from the ER (46). UGCG expression levels are also not decreased by GLTP KO, ruling out a decrease in available synthase as a reason for the decreased amount of GlcCer in the cell. Furthermore, neither sphingomyelin nor GalCer levels are affected by the absence of GLTP (10), with sphingomyelin being synthesized at the *trans*-Golgi from a ceramide pool transported by CERT and GalCer being synthesized by ceramide galactosyltransferase at the ER lumen from the ER-native ceramide pool (47), with neither thus being reliant on COPII-vesicle-mediated ceramide transport. The lack of response in the levels of these lipids, contrasting with the significant depletion of GSLs deriving from COPII-vesicle-transported ceramide, further strengthens the interpretation that GLTP is a key factor in facilitating vesicle-mediated ER export.

The persistence of GSL levels in GLTP KO conditions may also be partly explained by other VAPA-interacting proteins, such as OSBP and ORPs, also having been implicated in facilitating ER export (43, 48). Additionally, the localization of LTPs like FAPP2 and CERT are altered in response to GLTP knockout. FAPP2 is a strong candidate for a compensatory role in ER-to-cis-Golgi transport. As a Golgi-targeted glycolipid transporter capable of membrane tubulation (49), it may help form ER–Golgi contact sites. CERT might also contribute to ceramide transfer to both the cis- and trans-Golgi. In the absence of GLTP, FAPP2 likely becomes the primary GlcCer transporter. If GlcCer synthesis shifts to the ER due to disrupted vesicle trafficking, FAPP2 may also mediate its export, ensuring its availability for downstream processes. This is supported by the observation that while GLTP or FAPP2 knockout alone is viable, their combined loss is lethal (35), suggesting that both are essential for maintaining GlcCer and GSL export pathways.

The specific mechanism through which the FFAT-mediated GLTP/VAPA interaction drives the ER-to-Golgi vesicle transport is so far unclear. Having previously demonstrated that ligand binding serves an inhibitory role for the GLTP/VAPA interaction (8), and having now demonstrated the necessity of the GLTP/VAPA interaction for the facilitation of COPII vesicle export from the ER, we suggest that GLTP maintains the GSL homeostasis of the cell via regulating COPII vesicle transport through a ligand-binding feedback loop. Higher levels of GSLs would result in more GLTP in the *holo*-state, i.e. with a bound GSL ligand, decreasing the occurrence of interactions between GLTP and VAPA; this would in turn decrease COPII vesicle export, diminishing ceramide transport to the *cis*-Golgi and decreasing the synthesis rate and amounts of synthesized GSLs. As a result of decreased GSL levels, an increase in *apo*-GLTP, i.e. without a bound GSL ligand, would follow, as would an increase in the incidence of GLTP/VAPA interactions, leading to an increase in ceramide transfer via COPII export and a corresponding gradual increase in GSL levels.

In conclusion, we propose that GLTP maintains the glycolipid homeostasis in the cell by regulating the COPII-vesicle-mediated export of ceramide from the ER to the *cis*-Golgi. This mechanism depends on the interaction between GLTP and VAPA at the ER membrane and which is reversibly inhibited by the binding of GSL ligands to GLTP. The retention of COPII coat proteins at the ERES, as well as the previously observed retention of vesicular cargo, during the absence of GLTP and during the expression of interaction-inhibited ΔFFAT-GLTP appear to indicate a requirement of GLTP/VAPA interaction for COPII vesicle export to occur. The changes in localization of the COPII vesicle assembly protein Sar1A because of GLTP KO and GLTP/VAPA interaction inhibition also allows us to tentatively further pinpoint the COPII secretion disruption to being a consequence of the failure of Sec12 to activate Sar1A. These factors corroborate previous hypotheses of GLTP acting as a regulator of GSL levels in the cell and simultaneously entails a major shift in our understanding of GLTP and its roles in the cell. Disruption of GLTP or its interaction with VAPA impairs COPII vesicle formation and cargo transport, indicating the necessity of this interaction for vesicle export. Additionally, the discovery of GLTP localization to the nucleus during DNA replication introduces an entirely new and unexpected dimension to GLTP’s *in vivo* function, warranting further investigation into its potential nuclear roles. Ideally, investigation would also be carried out in other cell lines which have been rendered GLTP deficient through CRISPR/Cas9 knockout, in order to determine whether the effects of GLTP on vesicle trafficking are universal or cell line dependent. However, we believe that the work carried out thus far already provides a robust indication of the potential *in vivo* role of GLTP, as not only a glycosphingolipid transporter but as a regulator of glycosphingolipid homeostasis and vesicular trafficking as well.

## Experimental procedures

### Cell culture

HeLa cells were used in all experiments as described previously (10). The cells were validated and free from *mycoplasma* contamination. Cells were cultured at 37 °C in a 5% CO_2_ atmosphere. Growth medium was Dulbecco’s Modified Eagle’s Medium (DMEM), supplemented with 50 U/ml penicillin, 50 U/ml streptomycin, 4 mM L-glutamine, and 10 % by vol. fetal calf serum (FCS). Stable GLTP KO HeLa cell lines generated as described in (10) were used.

### Gene vector transfection for generation of GLTP recovery and mutant expressing cells

Plasmid DNA encoding for wild-type human GLTP (hGLTP) or mutant GLTP variants was transiently transfected into HeLa cells to study the recovery of GLTP function in GLTP-KO cells or the effects of mutations on GLTP function. Transfection was carried out through electroporation with a Neon transfection system (ThermoFisher Scientific, United States). Prior to transfection, cells were grown to confluency in 100 mm culture dishes, to a minimum of 5x10^6^ cells per plate. Cells were then harvested through trypsination and pelleted by centrifuging at 200 x *g* for 5 minutes; the pellets were resuspended in Neon resuspension buffer R (ThermoFisher Scientific, proprietary buffer for Neon transfection kit) and mixed with plasmid DNA (2 μg plasmid DNA per plasmid per 5x10^6^ cells). Cells were then electroporated at manufacturer’s recommended system settings for the Neon system for HeLa cells (1005 V, 35 ms pulse length, 2 pulses). After electroporation, transfected cells were resuspended in DMEM and plated onto new 100 mm culture dishes; cells were allowed to recover for 24 hours before being harvested for proteins or prepared for imaging. The expression vector used was pcDNA3.1(+), with plasmids individually expressing wild-type hGLTP and ΔFFAT-GLTP being used (10).

### Immunofluorescence, image analysis and quantification

Cells for immunofluorescence analysis were cultured as described above, with the addition of sterilized glass cover slips being added to the culture dishes prior to plating of cells. GLTP rescue cells or cells expressing GLTP mutants were transfected with the relevant plasmid DNA as described above before being plated for immunofluorescence preparation. Cells were grown until sufficient confluency (∼70%) had been reached, at which time the glass slides were retrieved from the dishes and prepared for imaging. Slides were washed with PBS to remove possible debris and then treated with 4% paraformaldehyde solution (diluted with PBS from 16% formaldehyde solution, ThermoFisher Scientific, United States) for 6 minutes to fixate the cells. Slides were then washed again and treated with a detergent solution (0.1% Triton X-100, MP Biomedicals Inc., United States) for 10 minutes to permeabilize cells; finally, slides were washed one more time and incubated overnight at 37 °C with a 5% BSA PBS solution to block nonspecific protein bindings. The following day, cells were stained with the relevant primary and secondary antibodies (1 h and 30 min incubations at 37 °C, respectively), with the glass slides then being mounted on object glasses (using ProLong Diamond Antifade mountant with DAPI, ThermoFisher Scientific, United States) before being imaged and photographed under a fluorescence microscope (Zeiss Axio Vert.A1, Germany). The polyclonal antibodies against human GLTP have previously been described and had an ELISA titer ≥ 1:100,000 against the designed antigen (8, 9, 50–52). The primary commercial antibodies used were α-Sec12 (rabbit, AB181212; Abcam, United States), α-Sec16 (rabbit, HPA005684; Sigma-Aldrich, United States), α-Sec23A (goat, PA5-19011; Invitrogen, United States), α-Sec23B (mouse, MA5-27262; Invitrogen, United States), α-Sec31A (rabbit, PA5-52147; Invitrogen, United States), α-UGCG (rabbit, bs-21562R; Bioss Antibodies, United States), α-Sar1A (rabbit, PA5-103872; Invitrogen, United States), α-CERT (rabbit, AB151285; Abcam, United States), α-FAPP2 (goat, AB38748; Abcam, United States), α-Golgin97 (mouse, A21270; Invitrogen, United States) α-glucosylceramide (rabbit, RAS_0010; Glycobiotech GmbH, Germany), α-lactosylceramide (mouse, MAS_0090; Glycobiotech GmbH, Germany), and α-galactosylceramide (rabbit, RAS_0030; Glycobiotech GmbH, Germany). The secondary antibodies used were AlexaFluor Plus 488 goat-anti-rabbit IgG (A32731, Invitrogen, United States), AlexaFluor Plus 488 goat-anti-mouse IgG (A32723, Invitrogen, United States), AlexaFluor 555 donkey-anti-goat IgG (A21432, Invitrogen, United States), AlexaFluor 555 goat-anti-mouse IgG (A21422, Invitrogen, United States), and AlexaFluor 555 goat-anti-rabbit IgG (A21428, Invitrogen, United States). Antibodies were validated by Western blotting to confirm specificity for the target protein indicated by a single band at the expected molecular weight in lysates from wild-type cells. Negative controls omitting the primary antibody were included to rule out non-specific binding of secondary antibodies. The ImageJ software (version 1.54i) was used for manual analysis of individual cell fluorescence intensity distribution in regions of interest (ROIs). The fluorescence intensity was measured in defined regions of interest (ROI) corresponding to ERES and the nuclear and perinuclear area and over the fluorescence intensity of the entire cell body. Data are presented as percent of intensity for the ROI ± SEM compared to the intensity for the whole cell body from at least 15 cells across independent experiments.

### Western Blot

Protein levels in the cells were analyzed through Western blotting. Cells were grown to confluency, harvested through trypsination and pelleted by centrifuging at 200x *g* for 5 minutes. The pellets were resuspended in Laemmli buffer and incubated at 95 °C for 3 x 5 minutes on a heating block to lyse cells. Protein concentration in the cell samples was determined by the Lowry method (53). Proteins in cell lysates were separated through SDS-PAGE (12% SDS gel) and transferred onto PVDF membrane (transfer run for 15 minutes at 45V and 45 minutes at 120V). Membranes were protein-blocked by incubation at RT for 1 h in 2.5% w/v powdered milk – PBS-Tween (PBST; PBS with 0.1% Tween) solution. Proteins were thereafter labeled through overnight incubation at 4 °C or 1-2 h at RT on rocking stirrer with 1:1000 dilute solution of primary antibody (in PBST stock solution with 0,5% BSA and 0,02% NaN_3_), followed by 1 h incubation at RT with 1:10 000 dilute solution of secondary HRP-conjugated antibody in milk-PBST solution. Protein bands were visualized through incubation of labeled membrane with Western Lightning Ultra enhanced chemiluminescent substrate solution (Revvity, United States) at RT for 10 minutes on stirring plate, followed by imaging on an iBright CL1500 Imaging System (Invitrogen, United States). Primary antibodies used were the same as for immunofluorescence as described previously, as well as α-GLTP (rabbit, Epo53223, custom antibody; Eurogentec, Belgium), α-VAPA (mouse, OTI1D3; Invitrogen, United States), and α-β-actin (rabbit, SAB5600204; Sigma-Aldrich, United States). Secondary antibodies used were goat-anti-rabbit HRP conjugated antibody (31460, Pierce Protein Biology/Invitrogen, United States), goat-anti-mouse HRP conjugated antibody (31430, Pierce Protein Biology/Invitrogen, United States), and rabbit-anti-goat HRP conjugated antibody (31402, Pierce Protein Biology/Invitrogen, United States). A Glutathione-S-Transferase (GST) pulldown assay was used with purified GST VAP fusion protein lacking the short VAP C-terminal transmembrane domain as described previously (8). Any bound GLTP were pulled down with glutathione beads that were washed several times. A Western blot analysis was performed afterwards to visualize the presence of GLTP using anti-GLTP antibodies, as well as anti-GST antibodies as controls.

### Synchronization and cell cycle analysis of HeLa cells

Cell synchronization assays were carried out to determine whether the intracellular localization and/or levels of GLTP were altered throughout the cell cycle. Double thymidine block was used to synchronize cells to the G1/S phase boundary; RO3306 block for the late G2 phase; and nocodazole block to arrest cells mid-mitosis (38, 39). For the double thymidine block, cells were grown to approximately 40%-50% confluency before culture media was removed and replaced with culture media containing 2 mM thymidine (Sigma-Aldrich, United States). Cells were then grown in thymidine-containing media for 18 hours at 37 °C, after which the thymidine-containing media was removed, the cells washed twice with PBS, and untreated DMEM added to temporarily end the thymidine block. Cells were incubated with untreated DMEM for 9 hours at 37 °C, after which the untreated media was again removed and replaced with another round of media containing 2 mM thymidine. Cells were incubated for another 18 hours at 37 °C in this media, before the thymidine-containing media was again removed, the cells washed twice with PBS, and untreated media added again to release the cells from the thymidine block. Cells were then harvested at different timepoints either for protein analysis or for immunofluorescence imaging.

For the RO3306 block, cells were grown to approx. 50% confluency before the culture media was removed and replaced with media containing 10 μM RO3306 (Sigma-Aldrich, United States). Cells were then incubated for 20 hours at 37 °C, after which the RO3306-containing media was removed, cells washed twice with PBS, and untreated DMEM added after which cells were harvested at different timepoints for protein analysis or immunofluorescence imaging.

For the nocodazole block, cells were grown to approx. 50% confluency before the culture media was removed and replaced with media containing 100 nM nocodazole (Sigma-Aldrich, United States). Cells were incubated with this media for 14 hours at 37 °C, after which the nocodazole-containing media was removed, cells washed twice with PBS, and untreated DMEM added after which cells were harvested at different timepoints for protein analysis or immunofluorescence imaging

### Flow cytometry – PI staining of the synchronized cells

Synchronized cells harvested at different time points were analyzed using flow cytometry to confirm whether their cell cycle stages corresponded with each harvest timepoint. For each timepoint, 500000 cells were centrifuged, and the growth medium was removed. The cell pellet was gently resuspended in 430 µL of ice-cold 1× PBS. 1 mL of 95% ethanol (-20 °C) was added dropwise with stirring, and the cells were stored at -20 °C for ≥12 hours prior to preparation for FACS analysis. Staining was performed 1 hour before FACS. The cells were centrifuged, the supernatant discarded, and the pellet resuspended in 500 µL of staining master mix (2% propidium iodide, 1% RNase, 0.005% Tween-20 in 1× PBS). The cells were then incubated in a water bath at 37 °C for 30 minutes, protected from light. Following incubation, the samples were placed on ice and kept in the dark until FACS analysis was conducted (BD Fortessa Jazz, Becton Dickinson, United States).

## Data availability

All data including imaging presented here are from at least three independent experiments. The datasets generated and analyzed during the current study are included in this published article and are available from the corresponding author on reasonable request.

## Statistical analysis of data

The data are presented as means ± SEMs for at least three independent experiments. The statistical significance was calculated using a Student’s t test for pairwise comparisons. The level of statistical significance was set at 0.05: *p ≤ 0.05, **p ≤ 0.01, and ***p ≤ 0.001.

## Acknowledgement

We would like to thank Pia Roos-Mattjus for her helpful comments and guidance.

## Abbreviations

The abbreviations used are:

ER: endoplasmic reticulum
ERES: endoplasmic reticulum exit sites
GLTP: glycolipid transfer protein
KO: knockout
GSLs: glycosphingolipids
VAP: vesicle-associated membrane protein
COPII: coat protein complex II
CERT: ceramide transport protein
OSBP: oxysterol-binding protein
ORP: OSBP-related protein
FAPP2: four-phosphate adaptor protein 2
UGCG: UDP-glucose ceramide glucosyltransferase
GlcCer: glucosylceramide
GalCer: galactosylceramide
LacCer: lactosylceramide
FFAT: diphenylalanine-in-an-acidic-tract;
LTP: lipid transfer protein

## Notes

### Competing Interest Statement

The authors have declared no competing interest.

### Summary of Updates

Figure 4 revised. Supporting information is added.

### Funding

- Jane and Aatos Erkko Foundation
- Magnus Ehrnrooth Foundation
- Svenska Kulturfonden
- Medicinska Understödsföreningen Liv och Hälsa
- Stiftelsen för Åbo Akademi
- Åbo Akademi University

