## Supplemental material for "Glycolipid transfer protein modulates vesicular trafficking from the endoplasmic reticulum in HeLa cells"

### Supplementary Figures

Figure S1. 3D surface plots of HeLa cells and quantification of fluorescence intensity for Sec23A.

Figure S2. 3D surface plots of HeLa cells and quantification of fluorescence intensity for UGCG, FAPP2 and CERT.

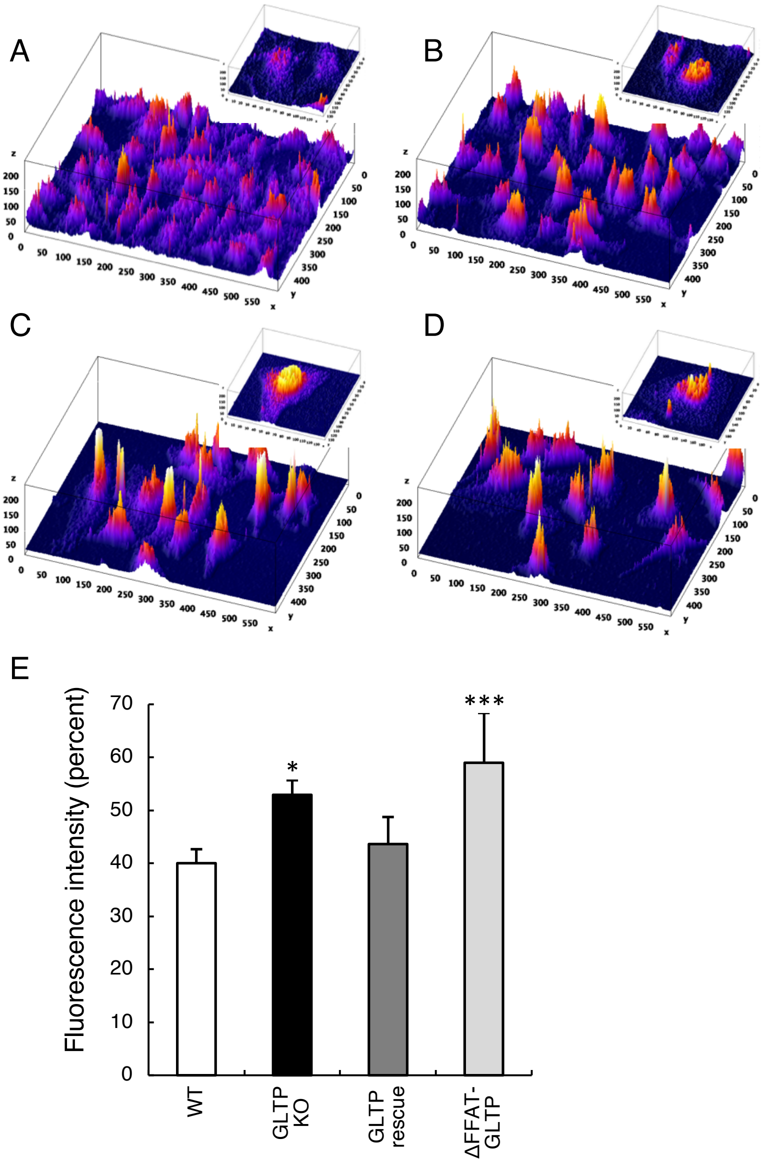


Figure S1. **3D surface plots of HeLa cells and quantification of fluorescence intensity for Sec23A**. We analysed the intensity of the fluorescence in HeLa cells with the ImageJ software using the 3D interactive surface plot to further analyze the changes in the intracellular localization of Sec23A as a function of GLTP knockout. We used ImageJ software to measure fluorescence intensity within specific regions of interest (ROI), including the nuclear, perinuclear, and ERES regions, and compared these values to the overall intensity of the entire cell body. (A) Surface plot of the fluorescence intensity for the expression of Sec23A in WT HeLa cells and (B) the expression of Sec23A in GLTP KO HeLa cells. (C) & (D) shows the fluorescence intensity of Sec23A in GLTP rescue HeLa cells and ΔFFAT-GLTP expressing HeLa cells respectively. (E) The fluorescence intensity was measured in defined ROIs corresponding to ER exit sites and the nuclear and perinuclear area and compared to the fluorescence intensity of the entire cell body. Data are presented as percent of intensity for the ROI ± SEM from at least 15 cells across independent experiments. The significance in difference between the WT group was tested using a Student's t test: $*p < 0.05$, $***p < 0.001$. The subpanels are higher magnifications (2x) of representative cells.

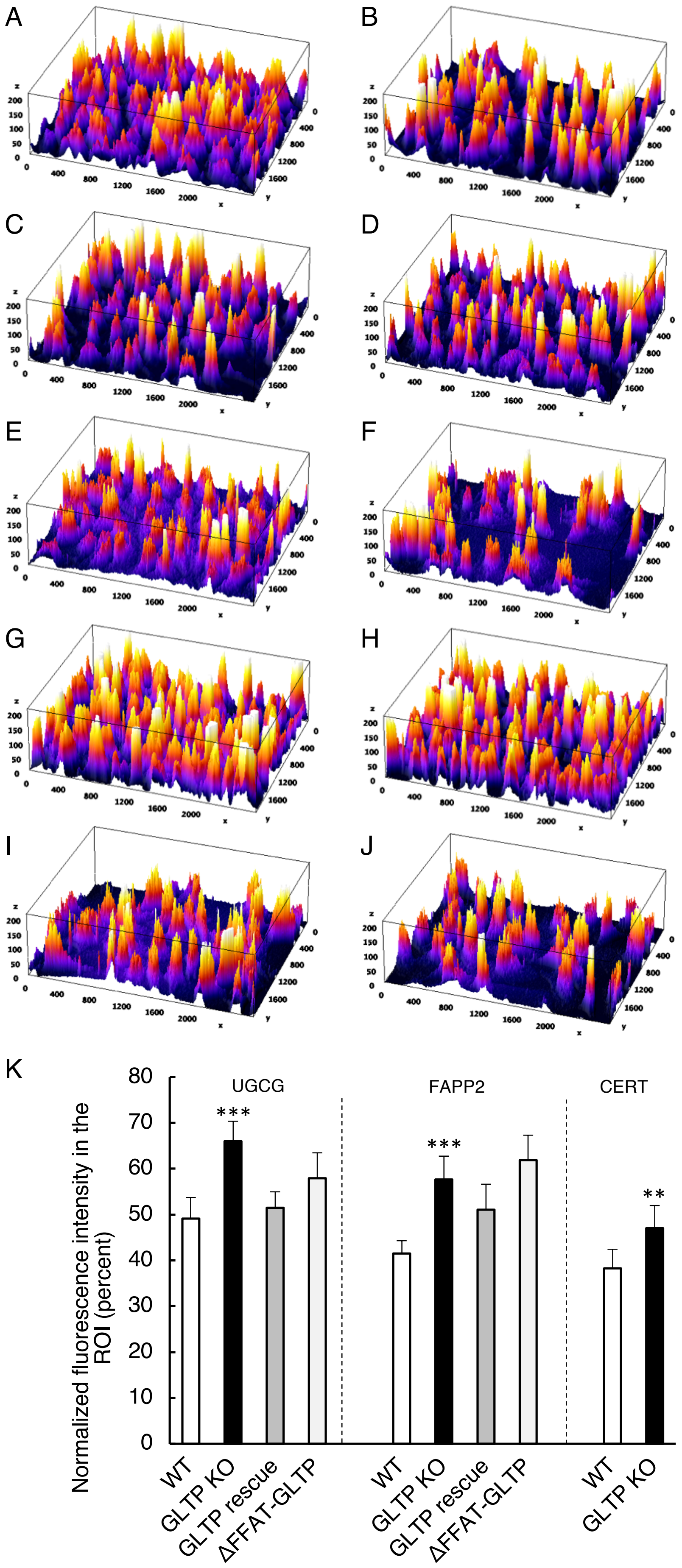
A
B
C
D
E
F
G
H
I
J
K
Normalized fluorescence intensity in the ROI (percent)
UGCG
FAPP2
CERT
***
***
**
WT
GLTP KO
GLTP rescue
ΔFFAT-GLTP
WT
GLTP KO
GLTP rescue
ΔFFAT-GLTP
WT
GLTP KO

Figure S2. **3D surface plots of HeLa cells and quantification of fluorescence intensity for UGCG, FAPP2 and CERT.** We analysed the intensity of the fluorescence in HeLa cells with the ImageJ software using the 3D interactive surface plot to further study the changes in the intracellular localization of different proteins as a function of GLTP knockout. We used ImageJ software to measure fluorescence intensity within specific regions of interest (ROI), including the nuclear, perinuclear, and ERES regions, and compared these values to the overall intensity of the entire cell body. (A) Surface plot of the fluorescence intensity for the expression of UGCG in WT HeLa cells and (B) in GLTP KO HeLa cells, (C) GLTP rescue HeLa cells and (D) ΔFFAT-GLTP expressing HeLa cells. (E) & (F) shows the fluorescence intensity for the FAPP2 protein expression in WT HeLa cells and in GLTP KO HeLa cells respectively and G) GLTP rescue HeLa cells and (H) ΔFFAT-GLTP expressing HeLa cells. (I) & (J) shows the fluorescence intensity for CERT protein expression in WT HeLa cells and in GLTP KO HeLa cells respectively. (K) The fluorescence intensity was measured in defined ROIs corresponding to ER exit sites, nuclear and perinuclear area over the fluorescence intensity of the entire cell body. Data are presented as percent of intensity for the ROI ± SEM from at least 15 cells across independent experiments. The significance in difference between the WT group was tested using a Student's t test: $**p < 0.01$, $***p < 0.001$.
